# Endothelial cells stably infected with recombinant Kaposi’s sarcoma-associated herpesvirus display distinct viscoelastic and morphological properties

**DOI:** 10.1101/2024.10.18.618233

**Authors:** Majahonkhe M Shabangu, Melissa J Blumenthal, Danielle T Sass, Dirk M Lang, Georgia Schafer, Thomas Franz

**Affiliations:** Biomedical Engineering Research Centre, Division of Biomedical Engineering, Department of Human Biology, Faculty of Health Sciences, University of Cape Town, Observatory 7925, South Africa; International Centre for Genetic Engineering and Biotechnology, Observatory 7925, South Africa; Institute of Infectious Disease and Molecular Medicine, Faculty of Health Sciences, University of Cape Town, Observatory 7925, South Africa; Division of Medical Biochemistry, Department of Integrative Biomedical Sciences, Faculty of Health Sciences, University of Cape Town, Observatory 7925, South Africa; Division of Physiological Sciences, Department of Human Biology, Faculty of Health Sciences, University of Cape Town, Observatory 7925, South Africa; Bioengineering Science Research Group, Faculty of Engineering and Physical Sciences, University of Southampton, Southampton, UK

**Keywords:** Cell mechanics, KSHV, mechanotyping, passive microrheology, vascular, lymphatic

## Abstract

Kaposi’s sarcoma-associated herpesvirus (KSHV) is a γ-herpesvirus that has a tropism for endothelial cells and leads to the development of Kaposi’s sarcoma, especially in people living with HIV. The present study aimed to quantify morphological and mechanical changes in endothelial cells after infection with KSHV to assess their potential as diagnostic and therapeutic markers. Vascular (HuARLT2) and lymphatic endothelial cells (LEC) were infected with recombinant KSHV (rKSHV) by spinoculation, establishing stable infections (HuARLT2-rKSHV and LEC-rKSHV). Cellular changes were assessed using mitochondria-tracking microrheology and morphometric analysis. rKSHV infection increased cellular deformability, indicated by higher mitochondrial mean squared displacement (MSD) for short lag times. Specifically, MSD at τ = 0.19 s was 49.4% and 42.2% higher in HuARLT2-rKSHV and LEC-rKSHV, respectively, compared to uninfected controls. There were 23.9% and 36.7% decreases in the MSD power law exponents for HuARLT2-rKSHV and LEC-rKSHV, respectively, indicating increased cytosolic viscosity associated with rKSHV infection. Infected cells displayed a marked spindloid phenotype with an increase in aspect ratio (29.7%) and decreases in roundness (26.1%) and circularity (25.7%) in HuARLT2-rKSHV, with similar changes observed in LEC-rKSHV. The quantification of distinct KSHV-induced morpho-mechanical changes in endothelial cells demonstrates the potential of these changes as diagnostic markers and therapeutic targets.

## 1. Introduction

The Kaposi’s sarcoma-associated herpesvirus (KSHV), also known as human gammaherpesvirus 8 (HHV-8), is the aetiological agent of Kaposi’s sarcoma (KS), a malignancy that affects the endothelium (Moore and Chang, 1995). KS presents primarily as cutaneous lesions composed of proliferative spindle cells of endothelial origin, often occurring in the immunocompromised such as people living with HIV (PLWH) and organ transplant recipients (Radu and Pantanowitz, 2013). KSHV infection in endothelial cells (ECs) can follow latent or lytic pathways. Latent infection, the default pathway, involves limited viral gene expression with the viral genome maintained as an episome in infected ECs. Spontaneous lytic reactivation in a subset of infected cells leads to extensive viral gene expression, new virion production, and infected cell death (Ganem, 2010). KSHV induces significant genetic and phenotypic changes in ECs, notably reorganisation of the cytoplasmic actin cytoskeleton by the viral latency-associated v-FLIP gene, which leads to the hallmark spindle-shaped cell morphology of KS-associated cells (Ganem, 2006).

Early diagnosis of KS is critical for a favourable prognosis. However, a confirmed KS diagnosis is resource-intensive and not always attainable in resource-limited settings that, for example, may have a shortage of experienced pathologists who can discern its various subtleties (Ackerman, 1979; Schneider and Dittmer, 2017). The resource-intensive diagnostic process involves histopathological identification of spindle cells from suspect lesion biopsies, which is usually confirmed by a positive latency-associated nuclear antigen (LANA) stain test. Although consistent use of combination antiretroviral therapy has led to regression of KS lesions in PLWH, poor outcomes persist in sub-Saharan Africa, exacerbated in part by high KSHV and HIV seroprevalence (Cesarman et al., 2019). The recent emergence of KS in virally suppressed PLWH underscores the need for further investigation of the pathobiology and treatment of KS (Palich et al., 2021).

During KSHV infection of target ECs, the cytoskeleton is implicated in viral entry and post-infection transcriptional dysregulation events, including endothelial-to-mesenchymal transition (EndMT) (Chandran, 2010; Gramolelli and Schulz, 2015). KSHV-induced actin reorganisation mediates the hallmark spindle morphology of KSHV-infected ECs. Given the central role of the actin cytoskeleton in modulating cell mechanics, the actin reorganisation indicates a possible association of KSHV infection with changes in intracellular viscoelastic properties (Fletcher and Mullins, 2010). Studies have demonstrated notable changes in cellular and organelle mechanical properties brought on by the infection of cells *in vitro* with herpesvirus and adenovirus (Andriasyan et al., 2021; Liu et al., 2020) and rubella virus (Kräter et al., 2018). If precisely defined, these intracellular stiffness properties can contribute to improved diagnosis and treatment of diseases associated with infections by these viruses. In the case of KS, the cell mechanical phenotype can serve as an alternative to identify and target the otherwise heterogeneous KS lesion cells.

The current study aimed to determine changes in the mechanical and morphological properties of ECs associated with their infection with KSHV as a precursor for KS tumorigenesis. The morphological and intracellular viscoelastic properties concomitant with latent recombinant rKSHV.219 infection of immortalised EC lines of vascular (HuARLT2) and lymphatic (LEC) origin were evaluated *in vitro*.

## 2. Materials and methods

### 2.1 Overview

The methodological approach used in the current study is illustrated in Figure 1 and described in the subsequent sections.

**Figure 1:**
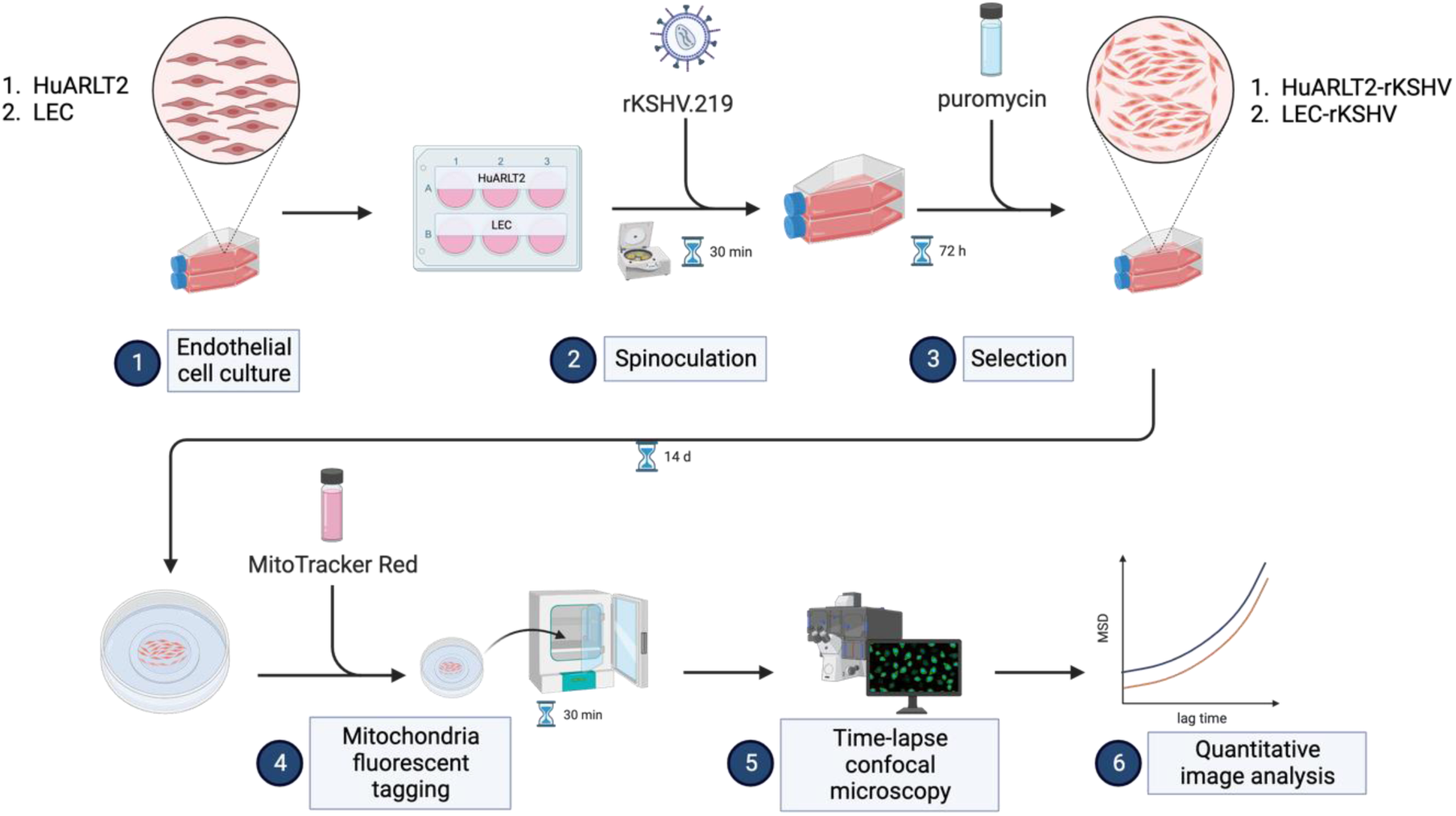
Methodology workflow. High-level illustration of sequential steps (1-6) of the approach used. (Created with BioRender.com)

### 2.2 Cell lines

BJAB-rKSHV.219 were established from BJAB cells (DSMZ No.: ACC 757), a human Burkitt lymphoma-derived cell line, via stable infection with a recombinant Kaposi’s sarcoma-associated herpesvirus (rKSHV.219). rKSHV.219, isolated from JSC-1 cells, expresses the green fluorescent protein (GFP) and the red fluorescent protein (RFP) driven by cellular EF-1α and KSHV lytic PAN promoters, respectively, and contains a gene for puromycin resistance as a selectable marker (Kati et al., 2015; Vieira and O’Hearn, 2004).

HuARLT2 cells are a conditionally-immortalised human umbilical vein endothelial cell line (HUVEC) derivative established via the expression of doxycycline-inducible transgenes simian virus 40 (SV40) large T antigen (TAg) and human telomerase reverse transcriptase (hTert) (May et al., 2010). LECs are a telomerase-immortalised human lymphatic endothelial cell (LEC) line (Bodnar et al., 1998; Hahn et al., 2012). Both HuARLT2 and LEC cell lines are well-established models for investigating KS mechanisms *in vitro* (Abere et al., 2017; Dubich et al., 2019). HEK-293T (DSMZ No.: ACC 305) cells are derived from human embryonic kidney 293 cells (HEK-293, Cellonex, C293-C) by stably expressing the transgenes SV40 and Tag. BJAB-rKSHV.219, HuARLT2 and HEK-293T cell lines were provided by Professor Thomas Schulz (MHH, Hannover, Germany). The LEC cell line was a gift from Dr Frank Neipel (Institute of Clinical and Molecular Virology, University Clinic, Erlangen, Germany).

### 2.3 Cell culture

BJAB-rKSHV.219 cells were maintained in Roswell Park Memorial Institute (RPMI; Sigma-Aldrich, South Africa) 1640 medium supplemented with 20% foetal bovine serum (FBS; Sigma-Aldrich) in the presence of 4.2 μg/ml puromycin (Abere et al., 2017).

HuARLT2 cells were grown and maintained in EGM-2MV BulletKit growth medium (Endothelial cell basal medium supplemented with SingleQuots; Lonza) at 37°C with 5% CO_2_ in the presence of 1 μg/ml doxycycline (Sigma-Aldrich).

LECs were maintained in EGM-2MV medium only.

HEK-293T cells were grown and maintained in Dulbecco’s Modified Eagle Medium (DMEM; Gibco) supplemented with 10% FBS.

For sub-culturing adherent cell lines and plating for downstream experiments, accutase (Biowest) was used to detach endothelial cell lines (HuARLT2 and LEC), and trypsin was used for HEK-293T cells. All cell quantifications were performed using the Countess Automated Cell Counter (Invitrogen).

### 2.4 Recombinant KSHV (rKSHV.219) production

For recombinant KSHV production, latently and stably infected BJAB cells (BJAB-rKSHV.219) were used as described previously (Kati et al., 2015). Briefly, BJAB-rKSHV.219 cells were cultured in 500 ml reusable spinner flasks (Beckman Coulter) at a density of 6 x 10^5^ cells/ml in 400 ml of glutamate-free RPMI supplemented with 20% FBS under constant agitation of 60 rpm for 4-5 days and in the presence of 2.5 μg/ml anti-human IgM antibody (Sigma-Aldrich) to induce the KSHV lytic cycle. After five days, the contents of the spinner flask were subjected to low-speed centrifugation in 50 ml Falcon tubes to remove cell debris and to collect supernatants in sterilised 250 ml centrifuge bottles (Corning). The virus was pelleted from supernatants via centrifugation at 14000 rpm for 6 hours at 4°C using a Beckman J2-21 high-speed ultracentrifuge fitted with a JA14 rotor (Beckman Coulter Inc., Fullerton, CA, USA). After carefully removing supernatants, virus pellets were re-suspended in 1 ml of EGM-2MV and stored at 4°C for use within four weeks.

### 2.5 Determination of rKSHV.219 titres

HEK-293T cells cultured in DMEM supplemented with 10% FBS and plated on a 96-well plate at a density of 3 x 10^5^ cells/well were infected with rKSHV.219 using the serial dilutions method and spinoculation, where plates were centrifuged at 450 x g for 30 mins at 30°C. Three days post-infection, GFP-positive cells in selected wells were manually counted using fluorescent microscope imaging and the virus titre was determined in infectious units per ml (IU/ml).

### 2.6 Infection of endothelial cell lines with rKSHV.219

HuARLT2s and LECs were infected with rKSHV.219 by spinoculation, as described before (Abere et al., 2017). Briefly, HuARLT2 cells and LECs were plated on a 6-well plate at a density of 5 x 10^5^ cells/well and subjected to rKSHV.219 infection via low-speed centrifugation of 450 x g for 30 mins at 30°C at multiplicities of infection (MOI) of 0.1 and 1, respectively, in the presence of 10 μg/ml polybrene (Sigma). EGM-2MV media containing 10 μg/ml polybrene was used for mock infections.

Inoculant media was replaced with normal EGM-2MV 8 hours after spinoculation. After three days, normal media was replaced with selection media comprising EGM-2MV media containing 5 μg/ml puromycin and 1 μg/ml doxycycline for infected HuARLT2 cells and 0.25 μg/ml puromycin only for infected LECs. Infected cells were maintained under selection for at least two weeks before particle tracking microrheology analysis. Latent infection in HuARLT2-rKSHV.219 and LEC-rKSHV.219 was confirmed qualitatively through GFP expression observed using a confocal fluorescent microscope (ZEISS LSM 880, Germany), controlled by Zen Black software version 2.3 (Carl Zeiss) under the emission filter of 488 nm.

### 2.7 Mitochondria-tracking microrheology

Mitochondria-tracking microrheology (MTM) was used to mechanically characterise the intracellular environment of cells. Cells were seeded on pre-equilibrated, uncoated, 4-compartment glass-bottom viewing dishes (µ-Dish 35 mm Quad–Ibidi, GmbH, Gräfelfing, Germany) suitable for confocal microscopy at a seeding density of 1.2 x 10^3^ cells/ml and kept overnight at 37°C with 5% CO_2_. EGM-2MV media was replaced with 100 nM of fluorescent mitochondria dye (Mito Tracker Red FM) in EGM-2MV. Each quad was filled with up to 300 μl of staining media, and the viewing dish was incubated away from light at 37°C with 5% CO_2_ for 30 mins. The staining medium was removed, and cells were rinsed with EGM-2MV twice before microscopic imaging.

All imaging was performed in a 37°C, 5% CO_2_ environmental chamber. For time-lapse confocal imaging of live cells, between eight and 12 non-overlapping fields of view (FOV) were acquired for each condition (i.e. viewing dish quad) using an inverted laser scanning confocal microscope equipped with a monochrome CCD camera (ZEISS LSM 880 with Airyscan, Germany) with a 63x oil immersion lens with 1.4 NA for 100 seconds at a frame rate of 10 Hz. An emission band pass filter of 585-640nm was used. To enhance the signal-to-noise ratio (SNR) and achieve a frame rate of at least 10 Hz, a 2×2 binning mode was used for a final spatial resolution of 0.33 μm/pixel. One thousand frames of 688 x 520 pixels were acquired for each FOV. At least two non-interacting cells from each FOV with a minimum of 50 identifiable punctate mitochondria were selected for particle tracking and analysis. Ten fluorescent images per experimental condition were collected using the same imaging set-up to identify latently infected GFP-expressing cells for particle tracking analysis.

Post-processing of time-lapse image series (x, y, t) was performed based on the approach described by Xu et al. (2018). Briefly, image stacks were pre-processed using FIJI (Schindelin et al., 2012) via the following pipeline: (i) cropping single cells (ii) conversion to 8-bit tiff stack (iii) histogram matching of all frames to the first frame to correct for bleaching (iv) background subtraction using a 40-pixel rolling radius (v) 2D deconvolution using 25 iterations to deblur the stack (vi) Gaussian blur filter with varying radii adjusted according to the SNR of each stack.

Thereafter, the plugin TrackMate (Tinevez et al., 2017) was used to segment image stacks, capture the spatial coordinates of near-punctate mitochondria, and generate and encode mitochondrial tracks into XML files. The Python-based Trackpy library version 0.5.0 (Allan et al., 2021) was used to process XML track files into mean squared displacement (MSD) curves and associated MSD scaling exponents (α-values). The MSD was reported for individual mitochondrion as time-averaged MSD (TAMSD), for multiple mitochondria of a single cell as ensemble-averaged MSD (EAMSD), and for multiple cells within the same group as the mean ensemble-averaged MSD (or bulk MSD). The EAMSD was determined according to:

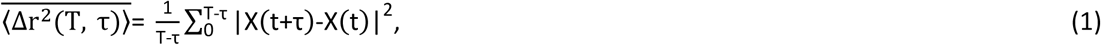

where Δr^2^ is the MSD, T is the total acquisition time, t is the iterative time point, τ is the lag time, ⟨ ⟩ represents the TAMSD, and — represents the EAMSD(Burov et al., 2013). The MSD varies nonlinearly with the lag time following a power law relationship for viscoelastic materials reminiscent of the intracellular environment, that is, ⟨Δr^2^(T, τ)⟩ ≃ τ^α^. The MSD power law exponent, α, is a derivative quantitative parameter based on the EAMSD. It describes the diffusive mode of tracked particles and is calculated as the logarithmic gradient of the MSD-τ curve (McGlynn et al., 2020):

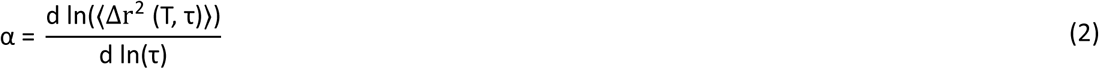

For extremely jagged EAMSD curves, indicative of highly dynamic mitochondrial fluctuations, the Savitzky-Golay filter with adaptive window size W_SG_ and polynomial order P_SG_ was applied. This adaptive approach ensured that the coefficient of determination (R^2^) for all power law fits within the lag time range of 0 s to 2 s was R^2^ ≥ 0.7.

### 2.8 Analysis of cell morphology

Phase contrast images of cells were captured using the Olympus IX81 inverted microscope (Olympus Corporation, Tokyo, Japan) at 10x magnification, controlled by Olympus CellSens Dimension v1.18 software (Olympus), for both HuARLT2 and LEC cell lines and their rKSHV-infected matches (Figure 2).

**Figure 2:**
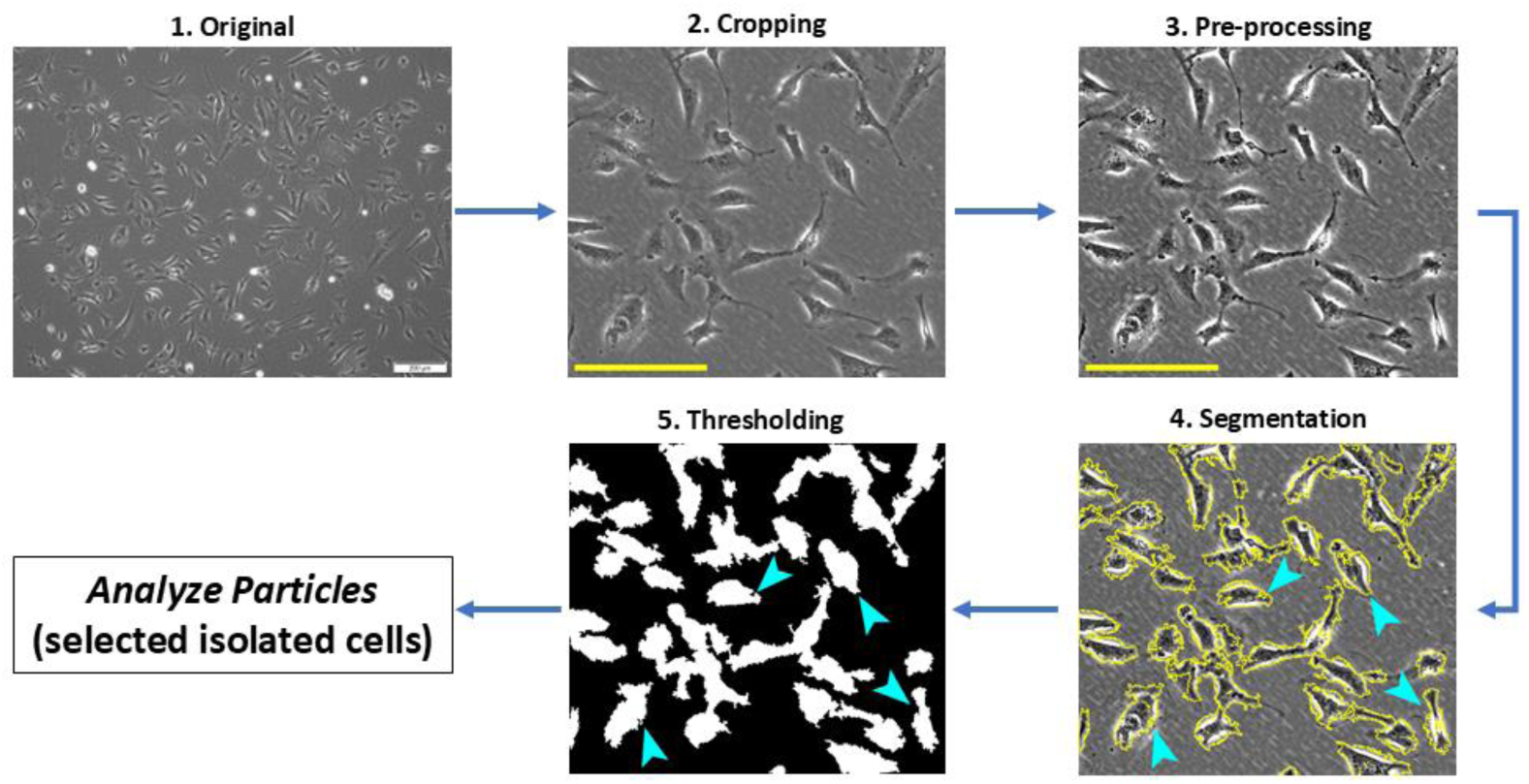
Image processing workflow for morphological analysis using FIJI and PHANTAST. Arrowheads indicate isolated cells earmarked for analysis.

Regions of interest (ROI) for analysis were extracted from phase contrast images and saved as separate images using FIJI. The confluency computation feature of the Phase Contrast Microscopy Segmentation Toolbox (PHANTAST) (Jaccard et al., 2014) plugin in FIJI was used to ensure that the extracted ROI images had comparable confluency levels.

The extracted ROI images underwent a series of pre-processing steps using FIJI. First, the images were converted to an 8-bit format to adjust brightness and contrast, enhancing cell boundary visibility. Afterwards, the images were converted to a 32-bit format to ensure compatibility with the PHANTAST plugin.

Image segmentation was performed using the PHANTAST plugin, and it involved adjusting the σ value, which controls the scale of Gaussian smoothing, and the ε value, which governs edge sensitivity or thresholding parameters for each image. Additionally, halo correction was selectively applied to enhance poorly resolved images and improve segmentation accuracy.

PHANTAST was used to generate binary masks that delineated the contours of individual cells within the images. These binary masks were then used to quantify the following cell shape parameters associated with the degree of cellular spindling in FIJI.

**Aspect ratio (AR)** is the ratio of the longest dimension to the shortest dimension in each cell to evaluate the degree of elongation of the cell,

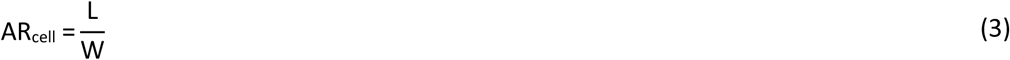

where L is the length of the longest dimension, and W is the length of the shortest dimension. Aspect ratio ranges from 0 to positive infinity and higher aspect ratio values indicate greater elongation and spindling.

**Roundness (R)** quantifies how closely the cell shape approximates a circle,

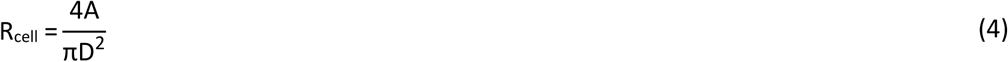

where *A* is the cell’s surface area, and *D* represents the major axis of the cell. Roundness ranges from 0 to 1, with higher values indicating a shape that is closer to a circle. However, it does not account for the cell boundary’s irregularities.

**Circularity (C)** quantifies how closely a cell’s perimeter resembles that of a circle,

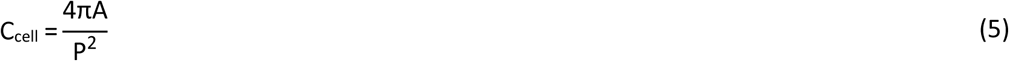

where A is the cell’s projected area, and P is the cell’s perimeter. Like roundness, circularity ranges from 0 to 1, with higher values indicating a more circular shape. However, unlike roundness, circularity considers the shape of the cell’s boundary.

### 2.9 Statistical analysis

The study aimed to determine cellular morphomechanical differences between uninfected vascular (HuARLT2) and lymphatic endothelial cells (LEC) and their rKSHV-infected counterparts. Statistical comparisons were performed between these biologically distinct groups.

The normality of the data was assessed using the Shapiro-Wilk test (α = 0.05). For normally or log-normally distributed data, Welch’s t-test was used to evaluate differences in intracellular viscoelasticity (bulk MSD and MSD power law exponent) and cell morphology (aspect ratio, roundness and circularity), with group statistics reported as mean ± SD. For non-normally distributed data, differences between groups were assessed using the Mann-Whitney U test, with group statistics reported as median and 95% confidence interval (Mdn [95% CI]).

Differences between groups were expressed as relative differences, with adjustments made for non-normally distributed data using Hodges-Lehmann differences. Statistical significance was considered at p < 0.05.

All experiments were conducted on two independent days, and statistical analyses were performed using Python 3.8 (Python Software Foundation, Python Language Reference, version 3.8) and GraphPad Prism 9.4.1 (GraphPad Software Inc., San Diego, CA, USA).

## 3. Results

The distinctive spindle phenotype is a hallmark signature in KSHV-infected primary endothelial cells. The present study quantified the morphological changes associated with spindling in latently rKSHV-infected HuARLT2 and LEC cells, as well as potential changes in intracellular mechanical properties for use as physical biomarkers. Stably KSHV-infected HuARLT2 and LEC cells were identified by GFP expression as a marker for rKSHV latent infection (see section 2.2). These cells and uninfected controls were fluorescently labelled with MitoTracker Red for particle tracking microrheology analysis (Figure 3).

**Figure 3:**
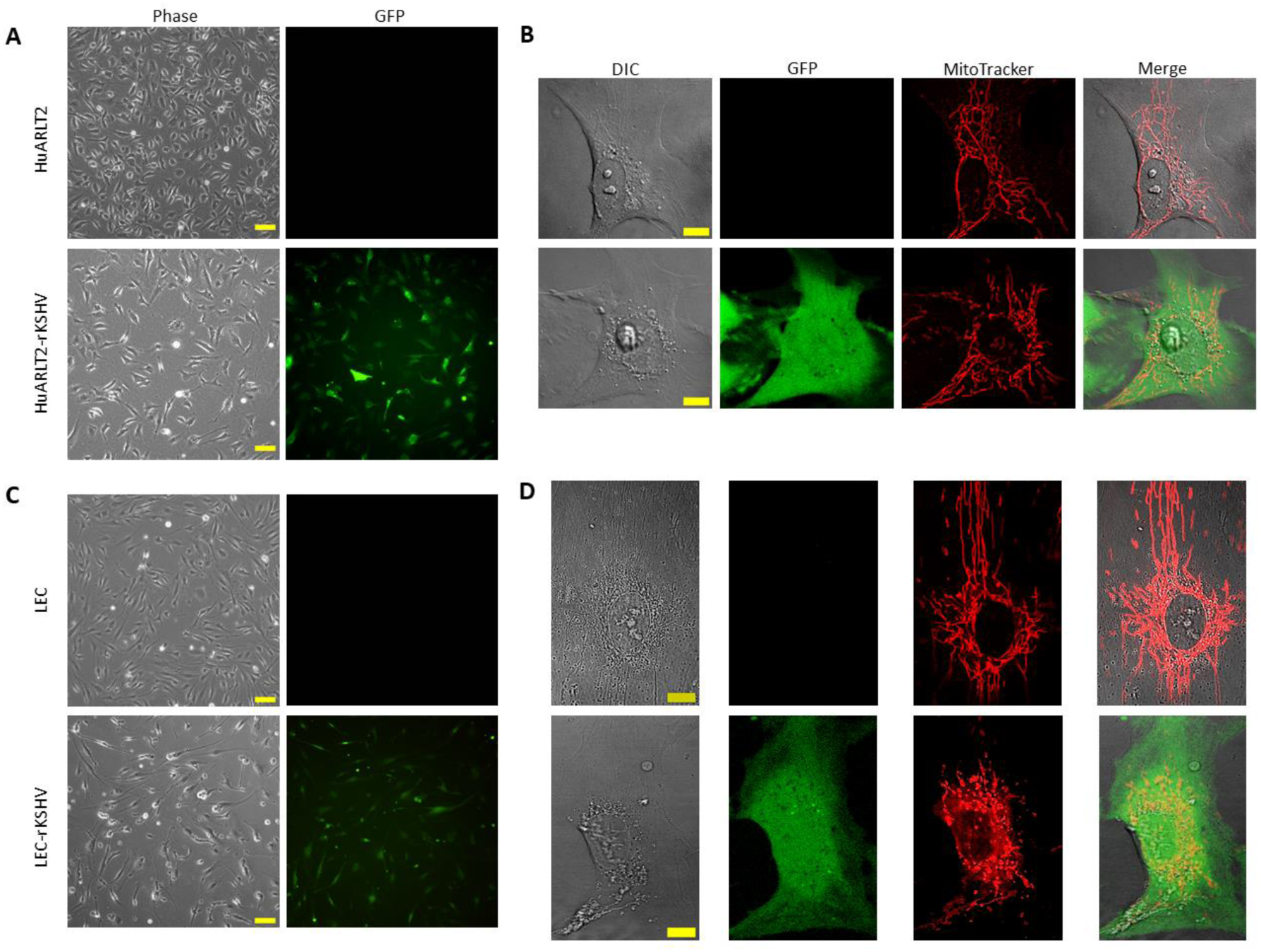
Representative micrographs of uninfected and latently infected (rKSHV) HuARLT2 and LEC cells. Phase contrast and fluorescence images of **(A)** HuARLT2 and HuARLT2-rKSHV cells and **(C)** LEC and LEC-rKSHV cells (10x, scale bar: 100 μm). Confocal images of single **(B)** HuARLT2 and HuARLT2-rKSHV cells and **(D)** LEC and LEC-rKSHV (60x, scale bar: 10 μm). GFP indicates latent rKSHV infection. MitoTracker Red fluorescently labels mitochondria for mitochondria-tracking microrheology.

### 3.1 rKSHV.219 establishes latency and induces morphological changes in HuARLT2 and LEC

Comparing HuARLT2-rKSHV and LEC-rKSHV with their uninfected controls, cell morphology analyses revealed considerably higher elongation in infected cells as quantified by cell aspect ratio (A), roundness (R), and circularity (C) (Figure 4A). HuARLT2-rKSHV exhibit a significantly higher aspect ratio (Mdn 2.315 [1.879, 3.532] vs 1.825 [1.306, 2.517], p = 0.0271), lower roundness (Mdn 0.432 [0.283, 0.532] vs 0.548 [0.397, 0.765], p = 0.0271), and lower circularity (Mdn 0.310 [0.276, 0.432] vs 0.472 [0.380, 0.557], p = 0.0022) than HuARLT2. Similarly, LEC-rKSHV display an increased aspect ratio (4.605 ± 2.914 vs 3.600 ± 1.396, p = 0.106), decreased roundness (0.257 ± 0.136 vs 0.308 ± 0.110, p = 0.152), and decreased circularity (0.256 ± 0.115 vs 0.352 ± 0.116, p = 0.0058) than the uninfected LEC, although only the change in circularity was significant.

**Figure 4:**
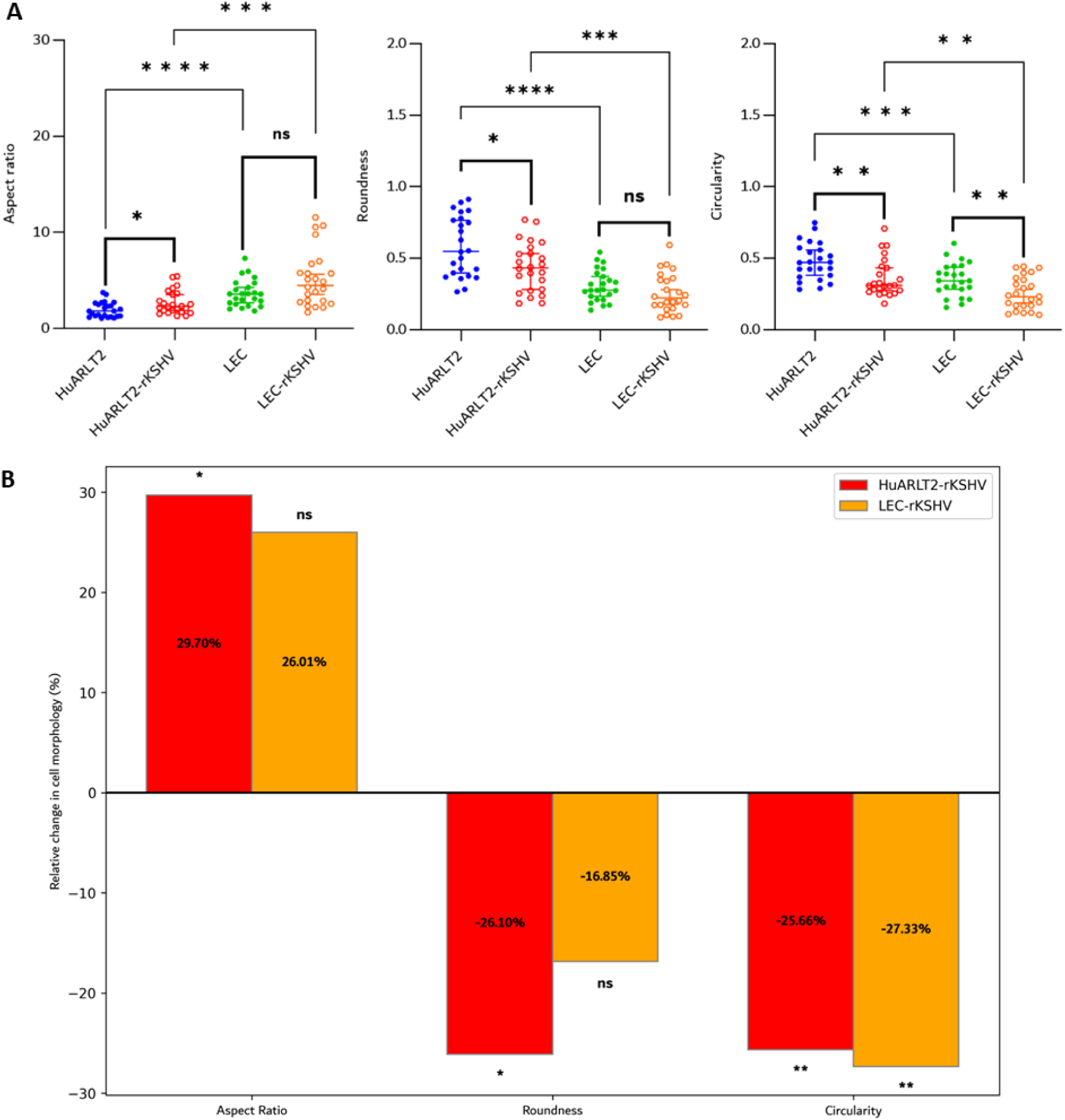
Cellular morphology. **A)** HuARLT2-rKSHV exhibit a significantly higher aspect ratio, lower roundness, and lower circularity than uninfected HuARLT2. Similarly, LEC-rKSHV display increased aspect ratio, lower roundness, and significantly lower circularity compared to uninfected LEC. LEC display stronger spindloid features than HuARLT2 both in the uninfected and the infected state. (N = 24 for each case. Plots represent mean ± standard deviation for normally distributed data and median and 95% confidence interval for not normally distributed data. * p < 0.05, ** p < 0.01, *** p < 0.001, **** p < 0.0001, ns non-significant.) **B)** Relative change in cell morphology characteristics (aspect ratio, roundness, circularity) for HuARLT2-rKSHV and LEC-rKSHV compared to the respective uninfected controls. (* p < 0.05, **p < 0.01, ns non-significant.)

When comparing uninfected HuARLT2 and LEC cells, the aspect ratio was significantly higher in LEC (Mdn 1.825 [1.306, 2.517] vs. 3.600 [2.691, 4.281], p < 0.0001), whereas roundness (0.583 ± 0.210 vs. 0.308 ± 0.110, p < 0.0001) and circularity (0.477 ± 0.129 vs. 0.352 ± 0.116, p = 0.0009) were significantly lower in LEC cells. Similarly, when comparing HuARLT2-rKSHV to LEC-rKSHV, LEC-rKSHV displayed a greater aspect ratio (Mdn 2.315 [1.879, 3.532] vs. 4.605 [2.879, 5.794], p = 0.0002), along with significantly lower roundness (0.583 ± 0.210 vs. 0.308 ± 0.110, p = 0.0002) and circularity (0.310 [0.276, 0.432] vs. 0.237 [0.169, 0.349], p = 0.0073)(Figure 4A).

In summary, rKSHV-infected HuARLT2 cells showed a significant 29.7% increase in aspect ratio, along with a 26.1% reduction in roundness and a 25.7% decrease in circularity compared to uninfected controls. In contrast, rKSHV-infected LEC cells showed a smaller, statistically insignificant 26.0% increase in aspect ratio, a 16.9% decrease in roundness, and a significant 27.3% decrease in circularity compared to their uninfected counterparts (Figure 4B). Latent rKSHV infection increased the aspect ratio and decreased roundness and circularity for both HuARLT2 and LEC, although the changes for roundness and circularity are not significant for LEC.

### 3.2 Increased MSD amplitude and decreased MSD power law exponent are observed with rKSHV-induced cellular elongation

The latently infected cells portrayed significantly higher mitochondrial MSD amplitudes than uninfected cells, with differences decreasing with increasing lag time (Figure 5 A and C). For short lag times between τ = 0 s and τ ≍ 1 s with predominantly passive mitochondrial fluctuations, the MSD amplitude of HuARLT2-rKSHV was 49.4% higher (p = 0.0001) at τ = 0.19 s and 33.0% higher (p = 0.0046) at τ = 1.03 s than that of HuARLT2 (Figure 5B). The MSD amplitudes of LEC-rKSHV were between 42.2% (p = 0.0062, τ = 0.19 s) and 38.0% (p = 0.0115, τ = 1.03 s) higher than for LEC (Figure 5D).

**Figure 5:**
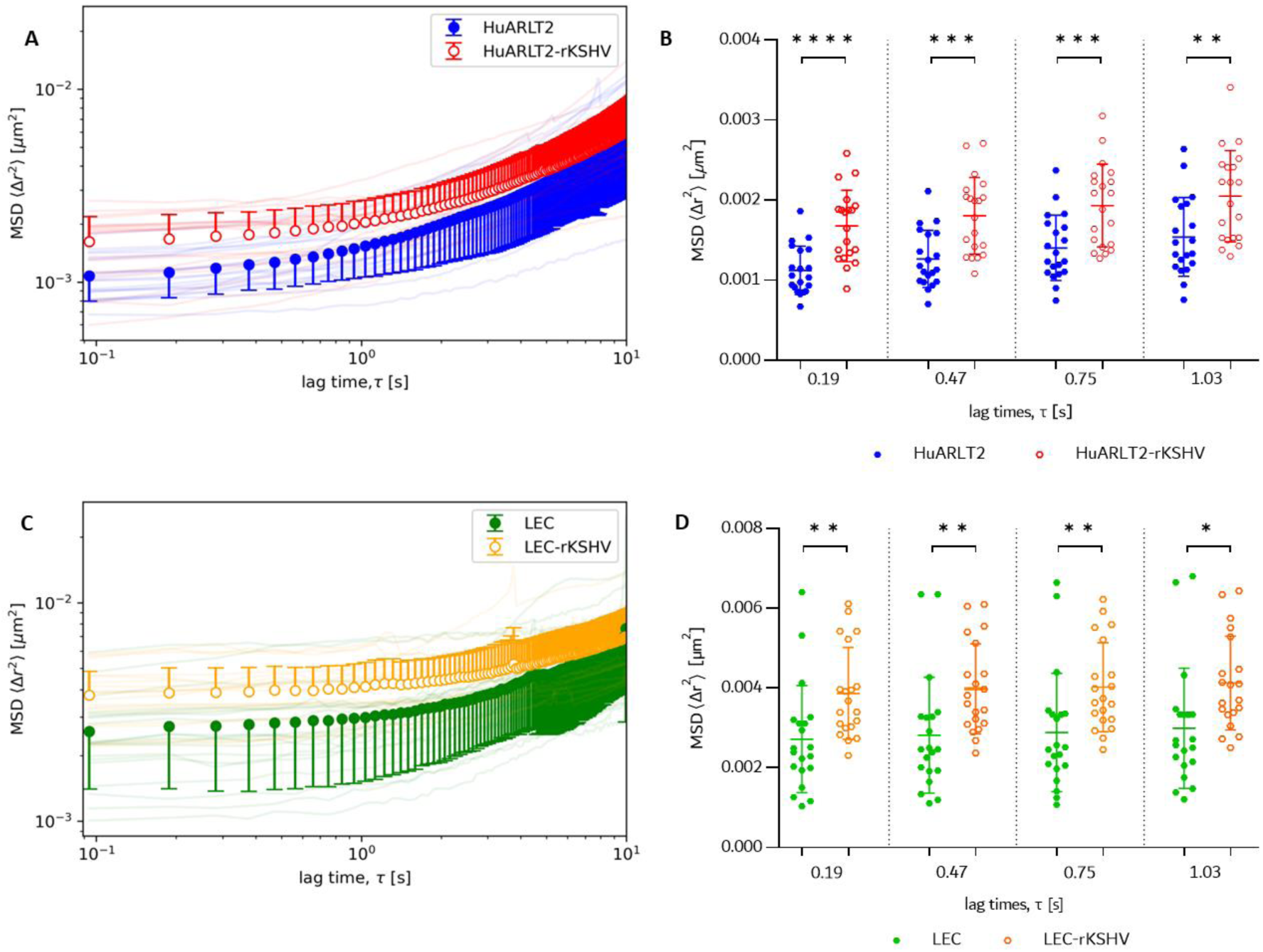
Mitochondria-tracking microrheology results of HuARLT2 (**A** & **B**) and LEC (**C** & **D**). **A** & **C**) Bulk MSD for uninfected (solid circles) and infected cells (open circles) for a lag time of 0 s ≤ τ ≤ 10 s. Error bars represent the standard deviation (SD). Ensemble-averaged MSD of individual cells are shown as faint lines. **B** & **D**) Ensemble-averaged MSD for individual cells and bulk MSD (mean ± SD) for lag times of 0 s ≤ τ < ≈1 s. * p < 0.05, ** p < 0.001, ***p < 0.0001, **** p < 0.00001 (N = 20).

When fitted to a power law function according to the behaviour of particles diffusing in a viscoelastic medium (Eqn.(2)), the infected cells (HuARLT2-rKSHV and LEC-rKSHV) exhibited significantly lower MSD power law exponent α than the uninfected counterparts (HuARLT2 and LEC) for a short lag time of 0 s ≤ τ ≤ 2 s (23.9%, p = 0.0395 and 36.7%, p = 0.0001, respectively) (Figure 6).

**Figure 6:**
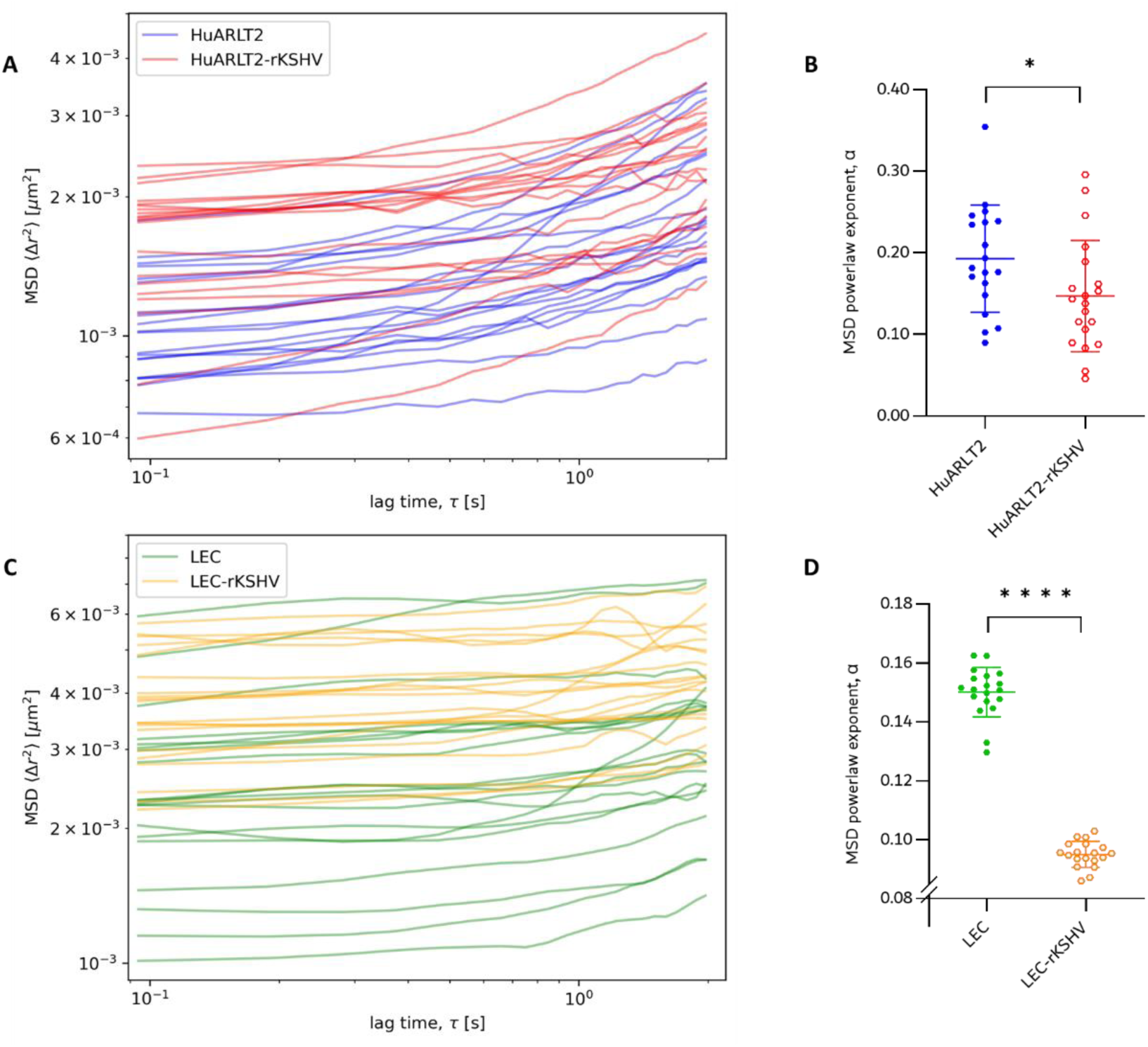
MSD and MSD power law exponent α HuARLT2 (**A** & **B**) and LEC (**C** & **D**) for a lag time of 0 s ≤ τ ≤ 2 s. Individual cell ensemble-averaged MSD curves (**A** & **C**) and fitted MSD power law exponent α (**B** & **D**) for uninfected (blue and green, solid markers) and infected cells (red and orange, open markers) for 0 s ≤ τ ≤ 2 s. The Savitzky-Golay filter was used with W_SG_ = 7 and P_SG_ = 3 to smoothen MSD curves for improved curve fitting for LEC-rKSHV and LEC (C). * p < 0.05, ** p < 0.001, ***p < 0.0001, **** p < 0.00001.

## 4. Discussion

The present study investigated changes in cellular morphology and viscoelasticity induced by rKSHV infection in HuARLT2 and LEC cells to identify potential morpho-mechanical markers of infection and early-stage KS tumorigenesis. The spindle phenotype in KSHV-infected endothelial cells is a crucial diagnostic parameter in KS histopathology; however, it has only been reported qualitatively to date (Matta et al., 2007).

Accurate histopathological diagnosis of KS spindle cells relies heavily on pathologist expertise, especially in the early stages of the disease, as other malignancies share a similar morphological profile (Schneider and Dittmer, 2017; Speicher et al., 2015). Identifying latently infected cells poses challenges due to the conservative gene expression profile of KSHV during latency (Ganem, 2006). Consequently, quantifying morphological changes is a key step towards robustly detecting infection in endothelial cells.

In the present study, relative to uninfected controls, HuARLT2-rKSHV cells displayed an approximately 1.3 times higher aspect ratio and about 0.74 times lower roundness and circularity, indicating marked transformation to a spindloid phenotype associated with rKSHV infection. The extent of spindling in LEC-rKSHV was lower than in HuARLT2-rKSHV, with an aspect ratio approximately 1.26 times greater than that of uninfected LEC and a reduction in roundness to about 0.83 times that of uninfected LEC. The reduced spindling was likely due to the higher pre-existing elongated morphology in uninfected LEC than in uninfected HuARLT2 (Figure 4 A), and therefore the aspect ratio and roundness may not be sensitive enough to confidently detect morphological changes associated with rKSHV infection. However, the circularity of rKSHV-infected LEC decreased to about 0.73 times to that of the uninfected control, supporting the presence of spindling (Figure 4 B). This quantification of spindling in rKSHV-infected HuARLT2 and LEC, known to be associated with actin cytoskeleton reorganisation, (Grossmann et al., 2006) represents a significant step in identifying EC morphological changes that are indicative of herpesvirus infection.

The MSD amplitude of tracer particles is directly proportional to the local intracellular deformability at a given lag time, τ (Hoffman et al., 2006). In the current study, HuARLT2-rKSHV and LEC-rKSHV showed 40.6% ± 7.0% (p < 0.0015) and 40.2% ± 1.9% (p < 0.0088) higher MSD amplitudes, respectively, than the uninfected controls for Brownian motion-dominated lag times of 0 < τ ≤ 1.03 seconds. This increase in MSD amplitude, therefore, suggests a decrease in local cellular stiffness, which coincides with the vFLIP-mediated spindling and altered actin cytoskeleton architecture. While the precise link between these MSD differences and their molecular aetiology remains unclear, the distinct upward shift in MSD amplitudes from uninfected to infected cells in both cell lines allows for effective distinguishing between infected and uninfected cell populations. Based on this result, rKSHV infection plausibly increases the mesh size between actin filaments, i.e. inter-filament distance, which then contributes to the more deformable phenotype observed in infected cells.

The mechanotyping of cancer cells can be inherently complex and depends on factors such as the specific malignancy and the analytical method and equipment used (Yu et al., 2022). Also, the MSD amplitude may be insufficient to characterise the viscoelasticity of the cytoplasm fully (Gal et al., 2013). However, in the current study, the MSD effectively captures the difference in deformability of the cells related to the presumed organisation of the actin cytoskeleton for infected HuARLT2 and LEC cells and their uninfected controls. The results reported here may be a unique phenomenon observed only within these specific cell lines, with the specific recombinant KSHV strain used, and at the specific time post-infection.

All values of the MSD power law exponent, α, across the cell lines were below 0.5, confirming the expected viscoelastic nature of the intracellular environment and constrained mitochondrial motion for both infected and uninfected cells (Wu et al., 2020). This indicates that mitochondrial fluctuations were predominantly passive and driven by thermal processes. The α value was 23.9% (p < 0.0395) and 36.7% (p < 0.0001) lower in the HuARLT2-rKSHV and LEC-rKSHV, respectively than in the associated uninfected controls for lag times of 0 < τ ≤ 2 seconds. The significant decrease in the power law exponent suggests a more viscous-like cytosol (Otten et al., 2012) in the infected than the uninfected cells. Despite higher MSD amplitudes, which suggest a larger actin cytoskeleton mesh size, the cytosol in infected cells is less permissive of mitochondrial fluctuations. The observed α values indicate alterations in the diffusive behaviour of mitochondria within the cells, and the decrease in α values may signify a shift towards more constrained or hindered mitochondrial motion, expected with an increase in cytosolic viscosity. The interpretation of the α value is multifaceted, influenced by factors such as cytoskeletal organisation, molecular interactions, and cellular microenvironment. Future studies could incorporate complementary techniques, such as active microrheology or fluorescence recovery after photobleaching (FRAP), to further validate the current findings.

The morphological and mechanical findings presented here suggest that the documented actin cytoskeleton reorganisation in endothelial cells during the early stages of KSHV infection (Ganem, 2010) is coupled with distinct and sustained changes in cellular morphology and viscoelasticity.

Interestingly, morphological changes of cells associated with spindling due to rKSHV infection were smaller in LEC than in HuARLT2 cells. In contrast, changes in mechanical properties (MSD and α) were the same in both cell lines if not higher in LEC than HuARLT2. This finding may suggest that intracellular mechanics is a more sensitive discriminator than cell morphology for detecting rKSHV infection in endothelial cell lines.

## 5. Conclusion

During KS tumorigenesis, the infection of endothelial cells by KSHV is deemed necessary, though insufficient to result in the malignant transformation of endothelial cells. While KSHV infection initiates the process of oncogenesis in endothelial cells, the development of KS only occurs when specific complementary conditions, such as immunosuppression and inflammation, are also present. It remains crucial, however, to detect KSHV infection as early as possible.

The current study demonstrates the cellular morphological and mechanical changes induced by rKSHV infection in HuARLT2s and LECs as a model for understanding the early phases of KS tumorigenesis. The results indicate that the cytoskeletal reorganisation known to result from KSHV infection of endothelial cells is associated with noticeable changes in cellular morphology and viscoelastic properties. While changes in cell morphology have been qualitatively observed through conventional histopathological staining in KS diagnosis, investigations into intracellular mechanics associated with this transformation have been lacking. The findings of the present study suggest that rKSHV infection leads to increased cell deformability, which may not only serve as a marker of early KS malignancy but also play a role in the pathogenesis of KS. Notably, increased cellular deformability following infection is observed in ECs of both vascular and lymphatic lineages implicated as primary targets of KSHV during KS tumorigenesis.

The present study underscores a biophysical approach’s significant role in elucidating virus-mediated oncogenesis. The findings contribute to understanding virus-mediated oncogenesis and developing novel strategies for early diagnosis of KSHV-associated malignancies. Augmentation with low-cost, automated, high-throughput microfluidics systems offers potential for point-of-care applications in under-resourced settings following the validation of physical cell markers of malignancy.

## Ethics

This work did not require ethical approval from a human subject or animal welfare committee.

## Data Availability

Data supporting this study are available on the University of Cape Town’s institutional data repository ZivaHub under the DOI https://doi.org/10.25375/uct.27043813 as Shabangu MM, Blumenthal MJ, Sass DT, Lang DM, Schafer G, Franz T. Data and software code for “Endothelial cells stably infected with recombinant Kaposi’s sarcoma-associated herpesvirus display distinct viscoelastic and morphological properties”. ZivaHub, 2024, DOI 10.25375/uct.27043813.

## Declaration of AI Use

No AI-assisted technologies were used in creating this article.

## Author Contributions

MM Shabangu: Conceptualization, Data Curation, Formal analysis, Investigation, Methodology, Project administration, Software, Visualization, Writing – Original Draft, Writing - Review & Editing

MJ Blumenthal: Investigation, Methodology, Writing - Review & Editing DT Sass: Formal analysis, Writing - Review & Editing

DM Lang: Methodology, Writing - Review & Editing

G Schafer: Funding acquisition, Methodology, Resources, Supervision, Writing - Review & Editing

T Franz: Conceptualization, Formal analysis, Funding acquisition, Resources, Supervision, Writing - Review & Editing

## Acknowledgements

We acknowledge the technical assistance of Ms Carla van Niekerk of the Confocal and Light Microscope Imaging Facility of the University of Cape Town.

